# Near real-time data on the human neutralizing antibody landscape to influenza virus to inform vaccine-strain selection in September 2025

**DOI:** 10.1101/2025.09.06.674661

**Authors:** Caroline Kikawa, John Huddleston, Andrea N. Loes, Sam A. Turner, Jover Lee, Ian G. Barr, Benjamin J. Cowling, Janet A. Englund, Alexander L. Greninger, Ruth Harvey, Hideki Hasegawa, Faith Ho, Kirsten Lacombe, Nancy H. L. Leung, Nicola S. Lewis, Heidi Peck, Shinji Watanabe, Derek J. Smith, Trevor Bedford, Jesse D. Bloom

## Abstract

The hemagglutinin of human influenza virus evolves rapidly to erode neutralizing antibody immunity. Twice per year, new vaccine strains are selected with the goal of providing maximum protection against the viruses that will be circulating when the vaccine is administered ∼8-12 months in the future. To help inform this selection, here we quantify how the antibodies in recently collected human sera neutralize viruses with hemagglutinins from contemporary influenza strains. Specifically, we use a high-throughput sequencing-based neutralization assay to measure how 188 human sera collected from Oct 2024 to April 2025 neutralize 140 viruses representative of the H3N2 and H1N1 strains circulating in humans as of the summer of 2025. This data set, which encompasses 26,148 neutralization titer measurements, provides a detailed portrait of the current human neutralizing antibody landscape to influenza A virus. The full data set and accompanying visualizations are available for use in vaccine development and viral forecasting.

## Introduction

The antigenic evolution of human influenza virus erodes the effectiveness of pre-existing immunity from prior infections and vaccinations, and is a major reason that the typical person is infected with influenza A virus roughly every five years^1,2^. The most rapidly evolving viral protein is hemagglutinin (HA); this protein is the major target of neutralizing antibodies, which are a strong correlate of protection against infection^3–5^. The HAs of human H3N2 and H1N1 influenza acquire an average of 3-4 and 2-3 amino-acid substitutions per year, respectively^6–8^.

Because of this rapid evolution, influenza vaccines are updated biannually with the aim of ensuring that they elicit neutralizing antibodies that protect well against current viral strains. Each update recommends a potentially new H3N2, H1N1, and influenza B strain(s) depending on if there is deemed to have been substantial antigenic change in circulating viruses relative to the strains in the prior vaccine. Recommendations are made twice per year: in September for the vaccine to be administered in the next Southern Hemisphere influenza season, and in February for the vaccine to be administered in the next Northern Hemisphere influenza season. Historically, recommendations were based primarily on antigenic measurements made using sera from previously naive ferrets infected with defined influenza strains^7,9^. However, there is growing recognition that the antibodies produced by humans with extensive lifetime exposure histories can differ from those produced by singly immunized ferrets^10–13^, so antigenic measurements made using human sera are now also increasingly considered^14–16^. A variety of evolutionary forecasting approaches are used to attempt to predict which strains might dominate in the coming season^17–20^. However, there remains a variable track record of choosing vaccine strains that turn out to be well matched to the strains that actually circulate the next season, and there is evidence that vaccine effectiveness is higher when the match is better^21–25^.

We recently developed a sequencing-based neutralization assay that enables simultaneous measurement of neutralization to many influenza strains^26,27^. The key innovation is to add unique nucleotide barcodes to each strain so that >100 viruses with different HAs can be pooled and assayed together, thereby enabling each column of a 96-well plate to measure the neutralization landscape of a serum against HAs from a wide diversity of strains. The neutralization titers measured using this sequencing-based assay are extremely similar to those measured using traditional one-virus versus one-serum neutralization assays^27^. We have shown that at least in 2023, the actual spread of different human H3N2 influenza strains in the human population was highly correlated with the fraction of individuals with low titers against those strains as measured by this assay^26^.

Here we designed a library of barcoded viruses that contained HAs from human H3N2 and H1N1 influenza strains that circulated as of April to May of 2025; this library continues to cover the HA diversity of these subtypes as of late summer 2025. We then used this library to measure neutralization titers for a set of 188 human sera collected from individuals of a wide range of ages at several geographic locations between October 2024 and April 2025. The resulting dataset of 26,148 neutralization titers provides a near real-time portrait of the human population’s neutralizing antibody landscape against influenza virus that can be used to help inform vaccine strain selection in September 2025 for influenza vaccines to be used in the 2026 Southern Hemisphere season.

## Results

### A library of influenza HAs that covers the antigenic diversity of human H3N2 and H1N1 influenza as of the summer of 2025

Our goal was to design a library of HAs that could be used to measure strain-specific neutralization titers prior to the September 2025 vaccine-strain selection (**Figure 1**). In April-May 2025, we chose a set of HAs to cover the genetic and antigenic diversity of human H3N2 and H1N1 strains that had been sequenced at that time. This library consisted of the HAs from 76 recently circulating human H3N2 strains and 38 recently circulating human H1N1 strains, as well as a total of 26 vaccine strains dating back to the 2014 vaccine for H3N2 and the 2010-2011 vaccine for H1N1 (**Figure 2**). We chose the recently circulating strains to include high-frequency HA protein genotypes in the 6-month period prior to library design, as well as strains observed within that time period that contained mutations at sites known to be antigenically important (see **Methods** for details).

**Figure 1.**
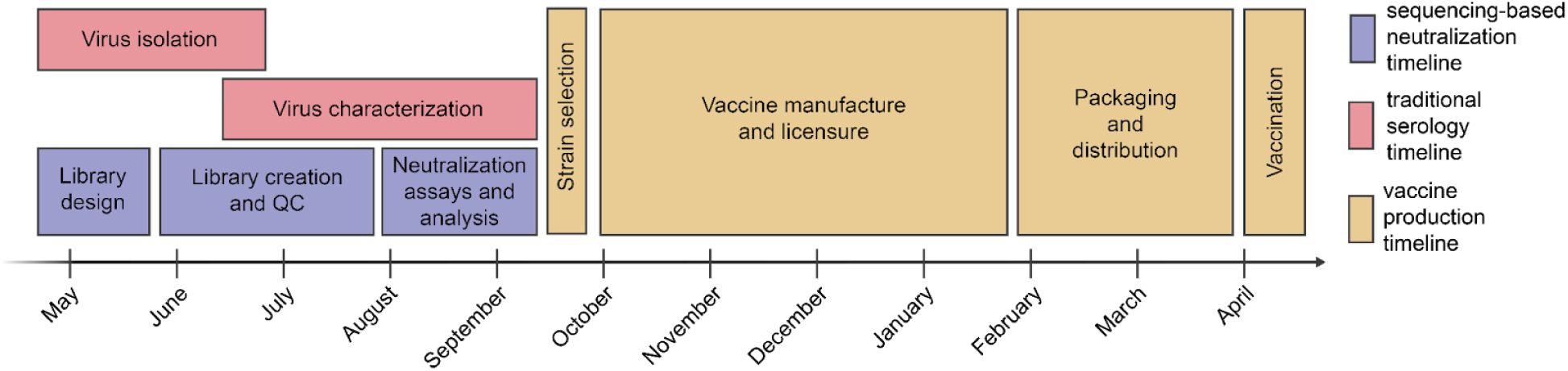
Typical timeline for data collection to inform influenza vaccine strain selection and subsequent production of vaccine for the Southern Hemisphere influenza season. Vaccine strain recommendations are made twice per year, once for the Southern Hemisphere vaccine and once for the Northern Hemisphere. In the timeline above typical for influenza vaccine strain selection and production for the Southern Hemisphere influenza seasons, shown in red and yellow are the periods for traditional virus isolation / characterization and vaccine production, respectively. The period shown in purple is for our sequencing-based neutralization assay measurements reported in the current study.

**Figure 2.**
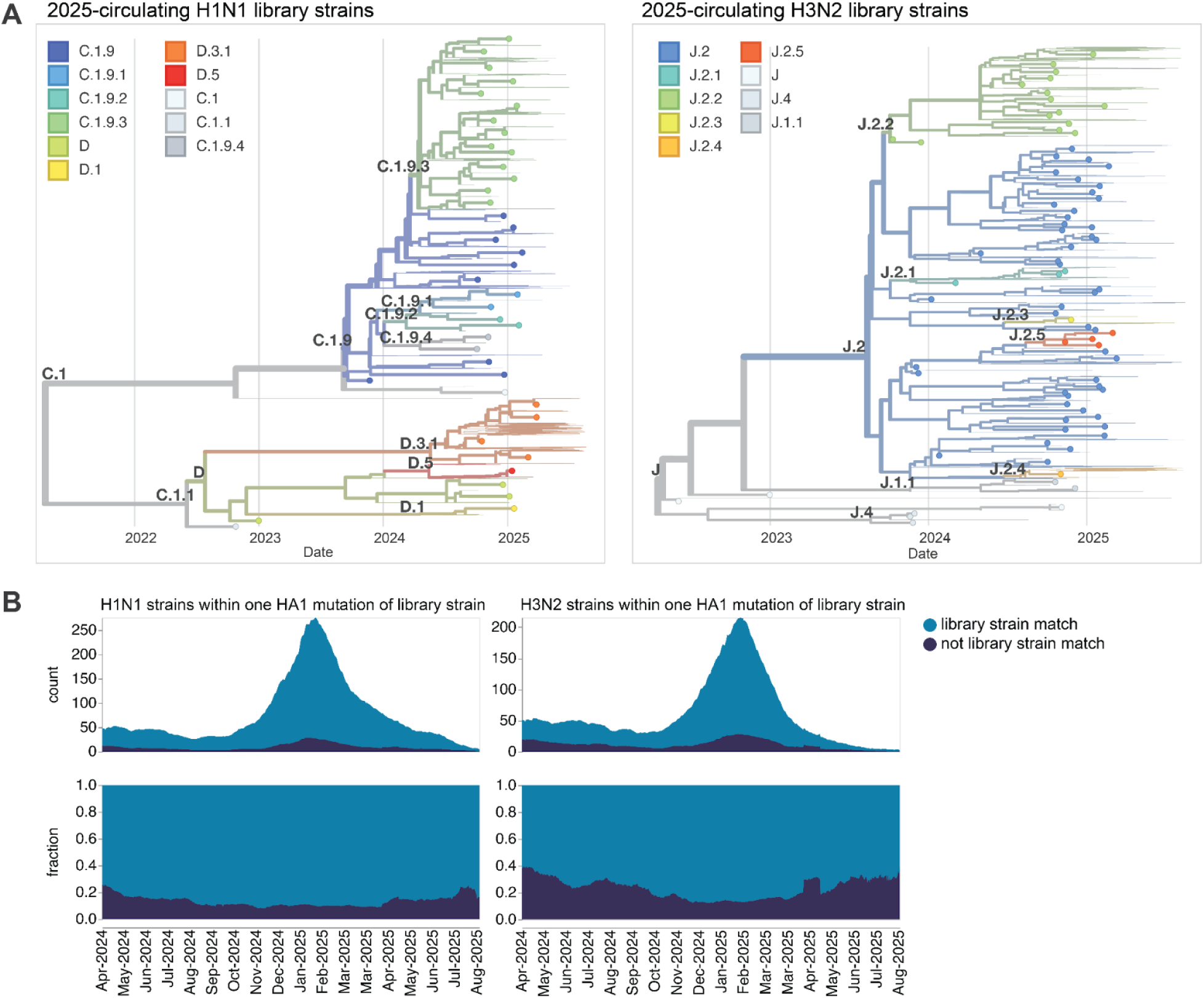
HAs included in the sequencing-based neutralization library. (**A**) Phylogenetic trees of HA genes of H1N1 and H3N2 strains included in the libraries. Strains included in the library are indicated with points, and other recent representative strains are shown in light lines. Colors indicate the subclade designation of each strain. These trees show only the recent strains designed to cover the current diversity of viral strains as well as the most recent vaccine strains (vaccine strains from 2024 seasons to present for both H1N1 and H3N2); note that the libraries also contained older vaccine strains dating back to 2012 for H3N2 and 2009 for H1N1. Interactive Nextstrain versions of these trees are available at (https://nextstrain.org/groups/blab/kikawa-seqneut-2025-VCM/h1n1pdm?f_kikawa=present_1&p=grid) and (https://nextstrain.org/groups/blab/kikawa-seqneut-2025-VCM/h3n2?f_kikawa=present_1&p=grid). (**B**) Count and fraction of all sequenced human seasonal H1N1 and H3N2 HAs available as of 2025-08-28 that are or are not within one HA1 amino-acid mutation of a strain in our libraries. The counts and fractions shown here are computed over a 10-day sliding window over the strain collection dates.

The HAs included in this library continue to effectively encompass most of the diversity of human H3N2 and H1N1 influenza as of late August 2025, with ∼78% of H1N1 and ∼67% of H3N2 viruses sequenced over the summer of 2025 being within a single HA1 amino-acid mutation of a strain in the library (**Figure 2**).

We individually generated barcoded viruses carrying each of the different HAs (**Figure 3**) using previously described approaches^26,27^. The HA genes were tagged with identifying 16-nucleotide barcodes and encoded the ectodomains from the naturally occurring recent H3N2 and H1N1 strains, and were incorporated into virions with the other seven viral genes derived from the lab-adapted A/WSN/1933 strain. We aimed to generate two or three distinct barcoded viruses for each HA variant to provide internal replicates in the sequencing-based neutralization assays. With these barcode replicates, after quality-control we had a total of 286 unique barcoded viral variants for the 140 different HAs. To create a library pool with all the variants at roughly equal titers of transcriptionally active particles, we pooled all the variants at equal volume, infected cells, extracted RNA, used barcode sequencing to quantify the relative transcriptional contribution of each variant, and then used these data to re-pool variants to balance the transcriptionally active particle titers for all HAs.

**Figure 3.**
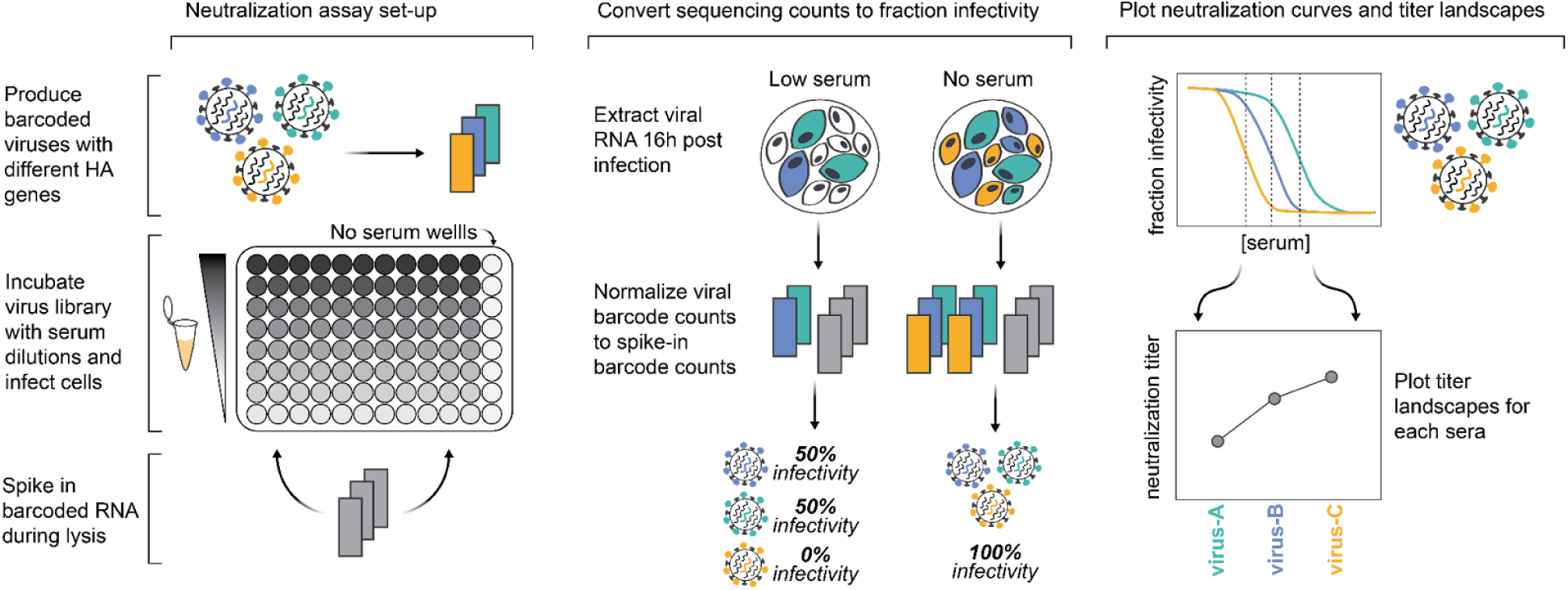
Sequencing-based neutralization assays enable rapid measurement of titers against many strains. Barcoded influenza viruses expressing different HAs are pooled and mixed with sera and MDCK-SIAT1 cells in an experimental set-up similar to traditional neutralization assays. To read out neutralization, viral RNA is extracted 16 hours post infection. A known concentration of barcoded RNA spike-in is added to each well and used to normalize viral RNA barcode counts, and these normalized counts are used to calculate the infectivity of each viral strain at each serum concentration relative to wells with no serum. Neutralization curves are fit to these percent infectivity values, and neutralization titers (defined as the reciprocal of the serum dilution at which 50% of viruses are neutralized) are calculated from these curves. The viral libraries used in this study contained viruses with 140 different HAs, and each plate was set up to assay 11 sera, meaning that each plate measured a total of 1,540 titers. Most HAs are represented multiple times in the viral libraries with several distinct barcodes, meaning most titers are measured in replicate in each plate.

### A diverse collection of sera collected from humans in late 2024 and early 2025

We assembled 188 human sera collected from individuals spanning from young children to elderly adults, and drawn from four sites around the world (**Figure 4**). These sera were collected between October 2024 and April 2025. Some sera are from individuals with some information on recent vaccination and infection status, for example in the EPI-HK study 19/42 (45%) participants reported receipt of the 2024/25 Northern Hemisphere influenza vaccine and 1/42 (2%) had PCR-confirmed influenza virus infection identified in-house within 182 days from date of serum collection (**Supplementary File 1**). However, many sera are residual samples from individuals with unknown infection and vaccination histories. Because the influenza immunity of most humans is due to a combination of infection and vaccination (with many individuals not receiving vaccines), and because infection often leaves a more durable imprint on antibody titers^28,29^ than vaccination, assaying residual sera from individuals with unknown vaccination histories as well as sera from well-characterized cohorts helps capture the diversity of anti-influenza serum antibodies across the human population.

**Figure 4.**
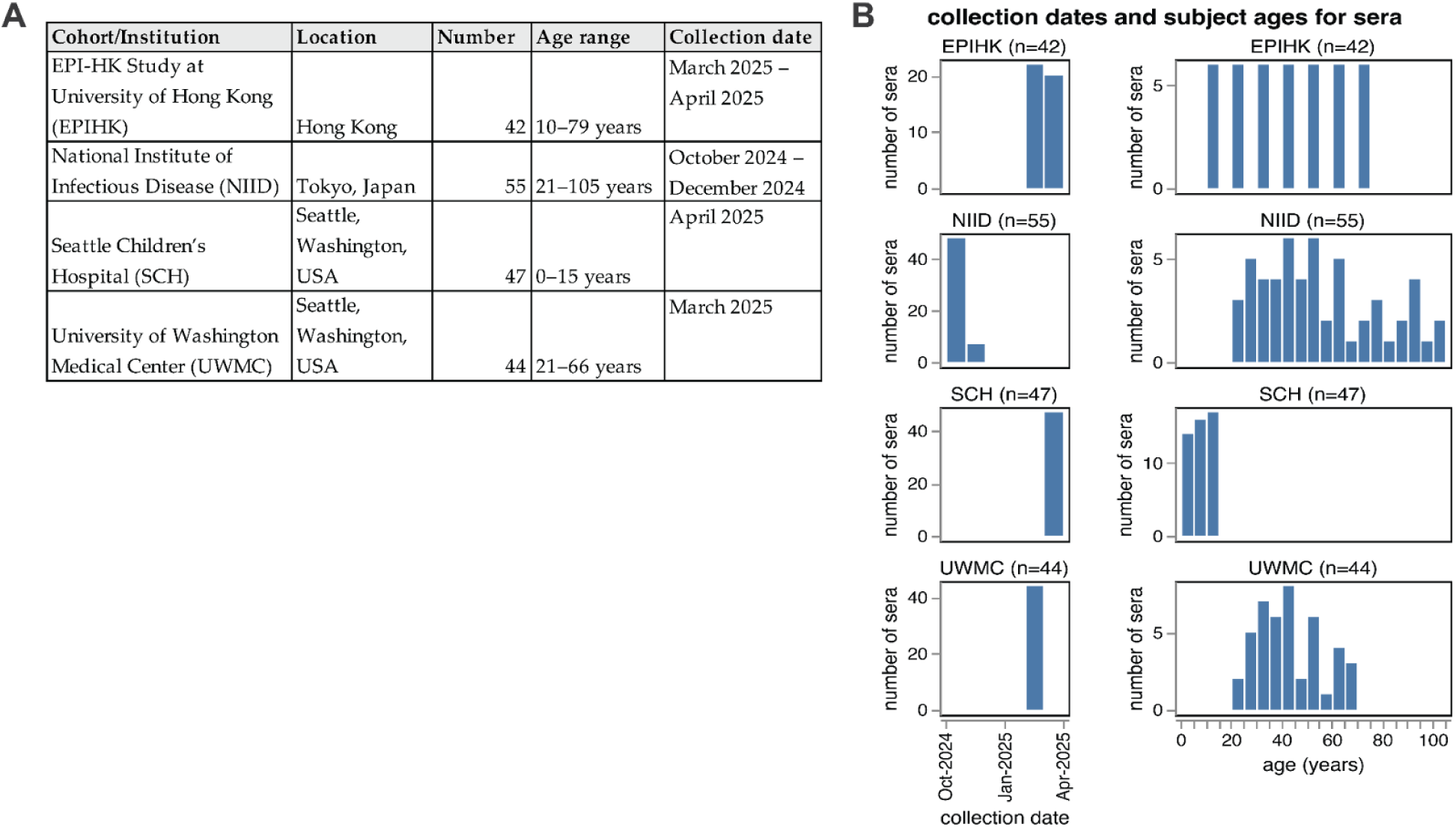
Human sera used in the neutralization assays. The human sera used in this study came from four different sources. (**A**) Details on the four groups of sera tested. All sera are from unique individuals with the exception of the NIID cohort; for that cohort 48 sera are from unique individuals pre-vaccination, and 7 sera are from 7 of those same individuals ∼1-2 months post-vaccination. (**B**) Distribution of collection dates and ages of individuals from which the sera was collected for each group.

### Human neutralizing antibody landscape against recent H3N2 and H1N1 strains

We used the sequencing-based neutralization assay (**Figure 3**) to measure neutralization curves for the 140 viruses against all 188 sera. After quality control to remove low-quality neutralization curves, we had a set of 26,148 neutralization titers (we quantify the neutralization titer as the reciprocal serum dilution that neutralizes 50% of the infectivity of a given viral strain as measured by our sequencing-based approach). Here we summarize major trends relevant to characterizing the neutralizing antibody landscape against the full set of viral strains across the tested sera; the **Methods** provide links to the full numerical titer data.

The neutralization titers were highly heterogeneous both across sera from different individuals and across viral strains (**Supplementary Figures S1-S3)**. Some of this heterogeneity was due to wide serum-to-serum variation in titers against all strains, as some individuals have generally higher anti-influenza neutralizing antibody titers than others. The heterogeneity in titers across sera was especially striking for children (eg, see the Seattle Children’s Hospital Cohort, SCH, in **Supplementary Figures S1-S3**), consistent with prior work^26,30^. To summarize the variation in titers due to differences between viral strains, we computed the median and interquartile range of the neutralizing titers against each viral strain across all sera (**Figures 5-7**).

**Figure 5.**
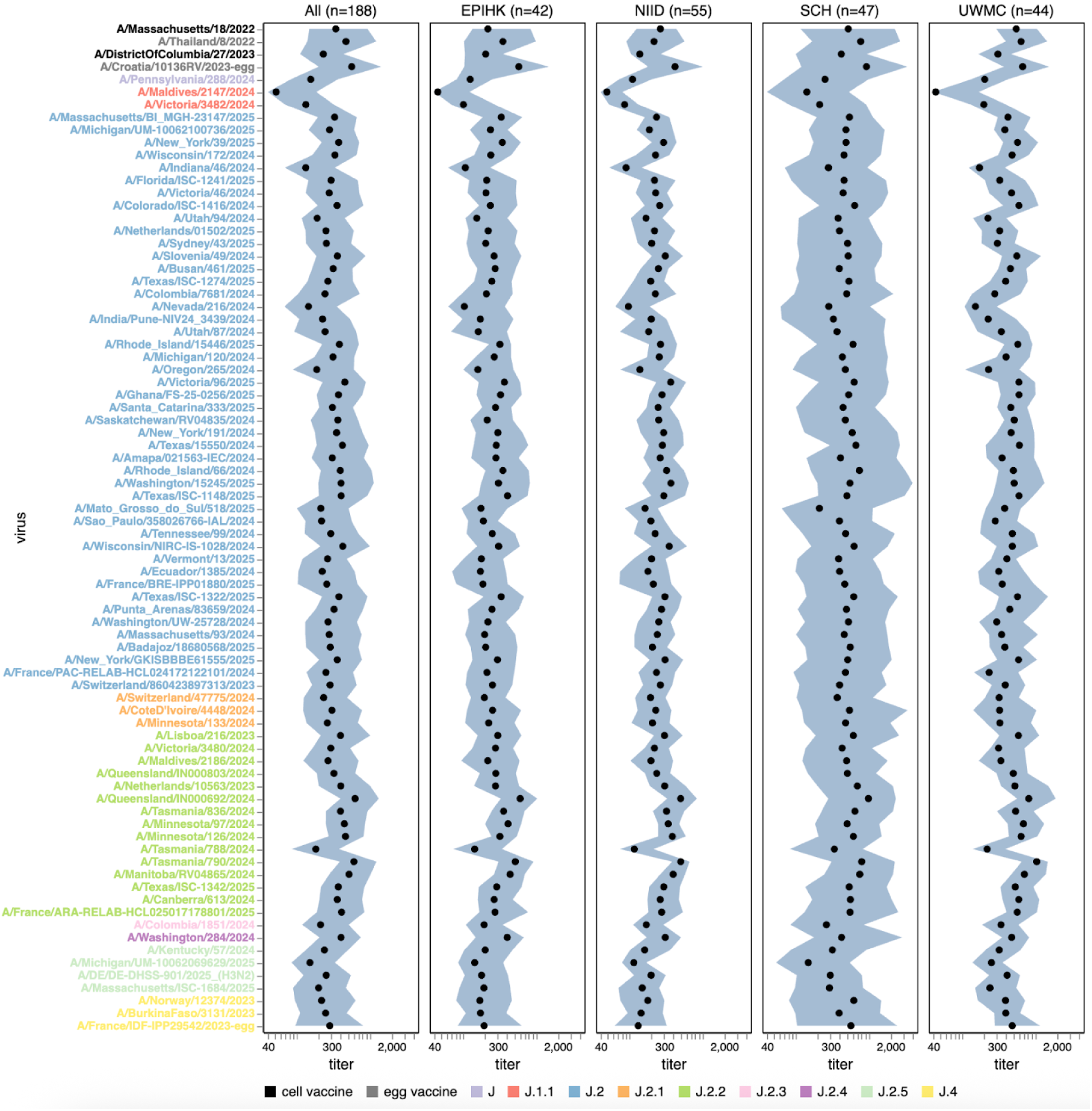
Human neutralizing antibody landscape against recent H3N2 strains. Median (black points) and interquartile range (shaded blue area) of the titers of the sera against the 76 H3N2 strains that capture current circulating diversity, as well as the last two sets of cell- and egg-produced vaccine strains. The leftmost plot shows the titers against all sera, and the remaining plots show titers for sera from each different cohort. Strain labels are colored by whether a strain is a cell- or egg-produced vaccine strain or its subclade per the legend at bottom. The lower limit of detection for titers in our neutralization assays was 40. See https://jbloomlab.github.io/flu-seqneut-2025/human_sera_titers_H3N2_recent_interquartile_range.html for an interactive version of this plot that allows mousing over points for details about individual viruses, and subsetting on sera from specific age ranges.

Among the recent H3N2 strains that capture current viral diversity, the median titers against different strains varied by more than five-fold (**Figure 5** and **Supplementary Figure S1**). The titers against the most recent (2025-2026) cell-produced H3N2 vaccine strain A/DistrictOfColumbia/27/2023 are comparable to those against many other strains in the library, suggesting that most currently circulating strains are not substantially antigenically advanced compared to this vaccine strain. However, some strains scattered across several subclades are neutralized by typical human sera substantially less well than this vaccine strain. The lowest median titers are to the J.1.1 subclade strain A/Maldives/2147/2024, and titers are also low to the related J.1.1 subclade strain A/Victoria/3482/2024; these strains share HA antigenic mutations I25V, S145N and I214T. However, the J.1.1 subclade has only been observed at low frequency recently, possibly suggesting other factors may limit its spread. Across multiple subclades, strains containing mutations at site 158 (eg, the J.2.5 strain A/Massachusetts/ISC-1684/2025) or both sites 158 and 189 (eg, the J.2.3 strain A/Colombia/1851/2024, the J.2 strain A/Mato_Grosso_do_Sul/18/2025, and the J.2.5 strain A/Michigan/UM-10062069269/2025) also tend to be neutralized poorly relative to other strains for some sera. Other strains that have relatively lower neutralization titers for some sera include the J.2.2 strain A/Tasmania/788/2024, and the J.2 strains A/Nevada/216/2024, A/Oregon/265/2024 and A/Indiana/46/2024. The same strains mentioned above with low median titers also tend to be ones with the highest fraction of sera with titers below a cutoff of 140 (**Supplementary Figure S4**), a feature that we previously showed correlated with strain evolutionary success in 2023^26^.

Among the recent H1N1 strains, the median titers were less variable across strains than for H3N2, with only about two-fold variation in median titers among strains (**Figure 6** and **Supplementary Figure S2**). The titers against the most recent (2025-2026) cell-produced H1N1 vaccine strain (A/Wisconsin/67/2022) were actually lower than those against most (but not all) other strains in the library. The titers against the most-recent egg-produced H1N1 vaccine strain (A/Victoria/4897/2022) were substantially higher than those against any recent circulating H1N1 strain, possibly because this strain contains several egg-adaptation mutations,^31,32^ including the R142K reversion and Q223R relative to the cell-produced vaccine strain^33,34^. Although no recent strains have a substantially lower median titer than others, there are a few strains in different subclades that do have substantial reductions in titer for subsets of sera, including the C.1.9.2 subclade strain A/KANAGAWA/AC2408/2025 and the C.1.9.3 subclade strain A/Ulsan/492/2025 (**Supplementary Figure S2)**. Both of these strains contain the G155E mutation, which can arise during lab-passaging of H1N1^35^ but is also sporadically observed in strains with no reported lab passaging, albeit only at low frequency among current strains. There was some modest variation across recent H1N1 strains in the fraction of sera that fall below a titer cutoff of 140, but this variation was much less than for recent H3N2 strains (compare **Supplementary Figures S4** and **S5**).

**Figure 6.**
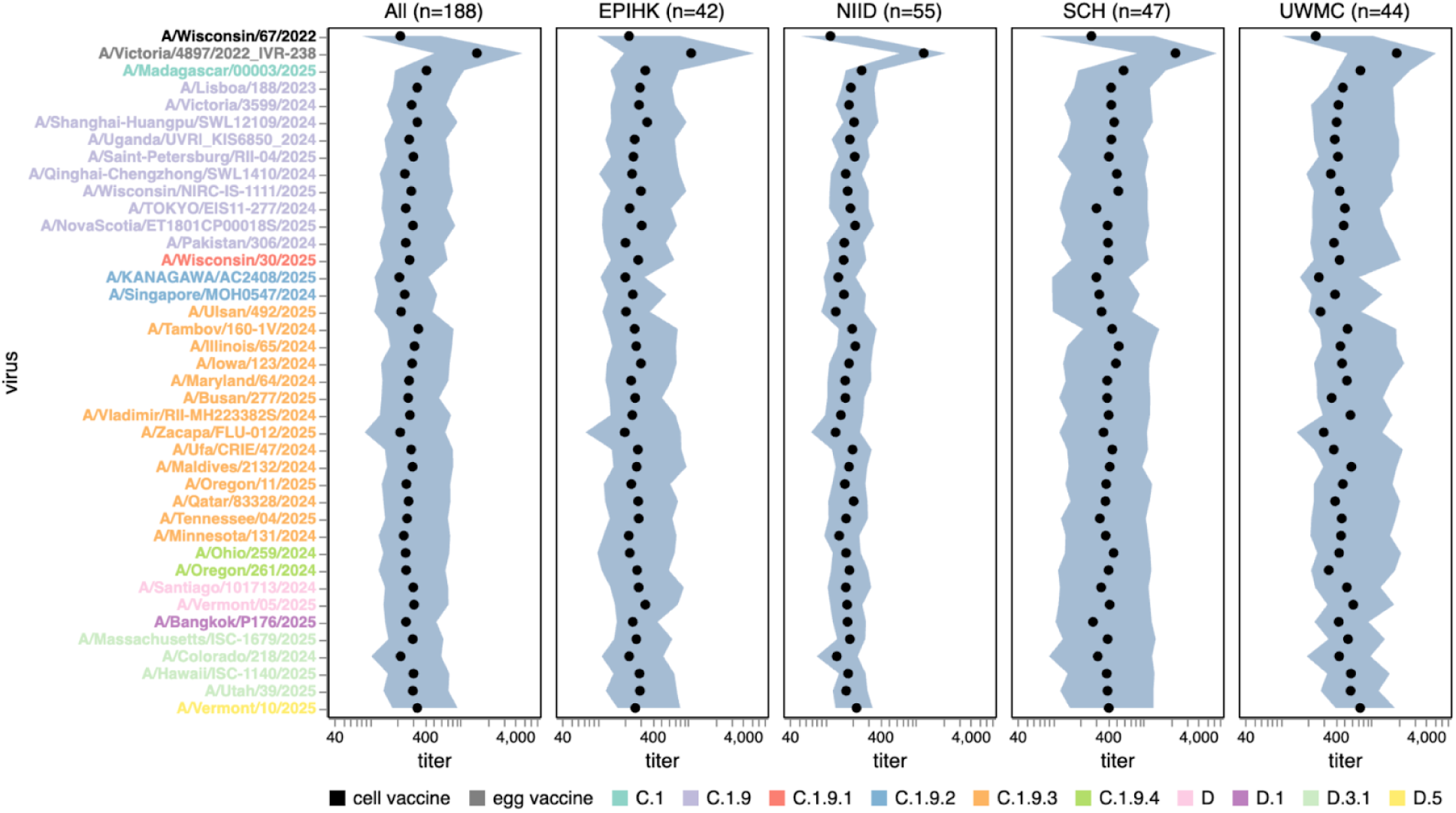
Human neutralizing antibody landscape against recent H1N1 strains. Median (black points) and interquartile range (shaded blue area) of the titers of the sera against the 38 H1N1 strains that capture current circulating diversity, as well as the last set of cell- and egg-produced vaccine strains. The leftmost plot shows the titers against all sera, and the remaining plots show titers for sera from each different cohort. Strain labels are colored by whether a strain is a cell- or egg-produced vaccine strain or its subclade per the legend at bottom. The lower limit of detection for titers in our neutralization assays was 40. See https://jbloomlab.github.io/flu-seqneut-2025/human_sera_titers_H1N1_recent_interquartile_range.html for an interactive version of this plot that allows mousing over points for details about individual viruses, and subsetting on sera from specific age ranges.

### Neutralizing antibody landscape to past vaccine strains

The library also contained past vaccine strains to complement the strains reflecting the current diversity of H3N2 and H1N1 seasonal influenza. Here we summarize notable trends in the titers of the human sera against these vaccine strains.

In recent years, separate strains have been chosen for vaccines produced in either cells or eggs, since human seasonal strains often do not grow well in eggs without adaptive mutations in HA^31,32^. The median titers to both H1N1 and H3N2 historical vaccine strains were consistently higher against the egg-produced vaccine strains relative to their cell-produced counterparts chosen for the same seasons, with titers to some pairs of egg-versus cell-produced vaccine strains for the same season differing by >5-fold (**Figure 7** and **Supplementary Figure S3**). This trend could be due to the antigenic effects of HA mutations selected by egg-passaging^31–33^ or reduced receptor avidity^36^ of egg-passaged viruses (the egg-produced viruses tended to be lower titer than the cell-produced equivalents on the MDCK-SIAT1 cells used in our experiments, favoring the latter hypothesis). For the H3N2 vaccine strains, the difference in titers between egg- and cell-passaged strains was lessened for more recent strains (**Figure 7** and **Supplementary Figure S3**).

**Figure 7.**
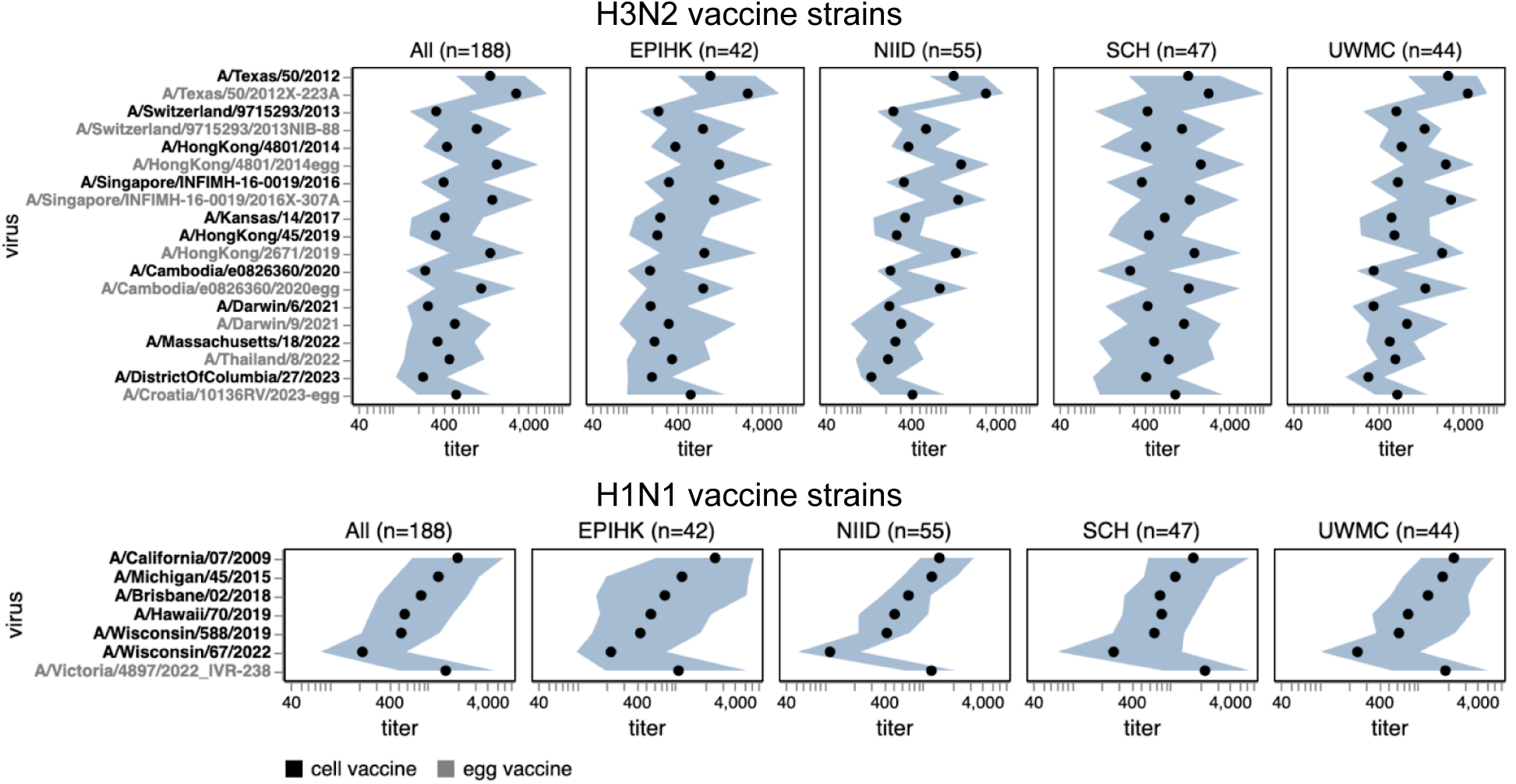
Human neutralizing antibody landscape against past vaccine strains. Median (black points) and interquartile range (shaded blue area) of the titers of the sera against past H3N2 (top) and H1N1 (bottom) vaccine strains; showing both the cell- and egg-produced vaccine strains for most seasons. The leftmost plot shows the titers against all sera, and the remaining plots show titers for sera from each different cohort. Strain labels are colored by whether a strain is a cell- or egg-produced vaccine strain per the legend at bottom. The lower limit of detection for titers in our neutralization assays was 40. See https://jbloomlab.github.io/flu-seqneut-2025/human_sera_titers_H3N2_vaccine_interquartile_range.html and https://jbloomlab.github.io/flu-seqneut-2025/human_sera_titers_H1N1_vaccine_interquartile_range.html for an interactive version of this plot that allows mousing over points for details about individual viruses, and subsetting on sera from specific age ranges.

Most sera had higher titers to older versus newer vaccine strains (**Figure 7** and **Supplementary Figure S3**), a result that makes sense as most sera in our study are from adults and extensive prior work has established that immune imprinting and back-boosting mean that humans tend to have higher titers to strains encountered earlier in their lives^1,12,37^. However, this trend is lessened or even reversed in the two cohorts that include children (SCH and EPI-HK), a fact that is most easily seen by using the sliders in the interactive versions of **Figure 7** and **Supplementary Figure S3** linked in their legends to subset just on sera from children—perhaps because the older vaccine strains only circulated before these children were born.

### Interactive visualization of neutralization titers in a phylogenetic context

We integrated our neutralization titer dataset into interactive visualizations of each subtype’s HA phylogeny using Nextstrain^38^ (**Figure 8** and links to interactive trees in figure legend). These trees show the phylogenetic relationships among HA sequences alongside a measurements panel where neutralization titers can be simultaneously visualized and compared to the tree^39^. The phylogeny and measurements panel are linked such that filtering or coloring the tree simultaneously filters or colors the titer measurements.

**Figure 8.**
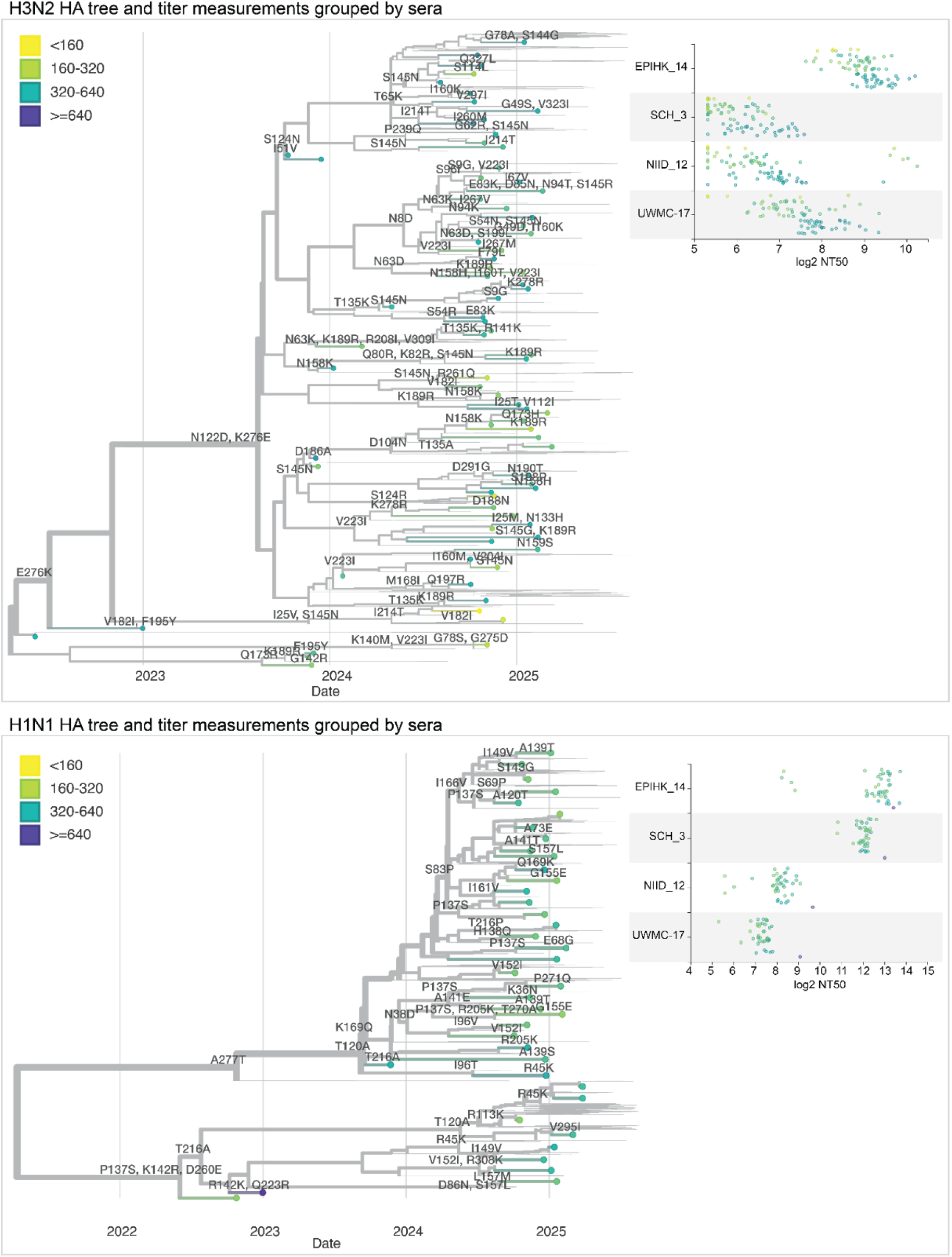
Static views of interactive Nextstrain trees showing the neutralization titers shown on HA phylogenetic trees. Phylogenetic trees of the HA gene of human H3N2 or H1N1 influenza, showing recent strains in the libraries as points, with thin lines representing other recent sequences. Circles representing the tips are colored by the median neutralization titer across all sera as measured in the current study. HA mutations are labeled on branches. The panels at right show the titers for four example sera across all viruses in the library. See https://nextstrain.org/groups/blab/kikawa-seqneut-2025-VCM/h3n2?c=median_titer&f_kikawa=present_1 and https://nextstrain.org/groups/blab/kikawa-seqneut-2025-VCM/h1n1pdm?c=median_titer&f_kikawa=present_1 for interactive Nextstrain versions of these nucleotide trees. Code to generate these nucleotide trees is available at https://github.com/blab/kikawa-seqneut-2025-VCM/. We also generated protein sequence-based trees (where branch lengths are in units of amino-acid mutations) for only the HAs in the sequencing-based neutralization assay libraries that can be viewed at https://nextstrain.org/community/jbloomlab/flu-seqneut-2025@main/H3N2 and https://nextstrain.org/community/jbloomlab/flu-seqneut-2025@main/H1N1.

We colored tree tips by median neutralization titers across all sera, annotated the tree with HA1 amino-acid mutations, and used the measurements panel to examine titers for specific viral strains and sera (**Figure 8**). For instance, the phylogeny shows how a subset of H3N2 strains with mutations 158K and 189R have lower median titers, further interactive examination of the tree shows those viruses are in the J.2.5 subclade **(Figure 8)**. The corresponding measurements panel shows how there is substantial variation across individual sera.

The interactive trees also make it possible to explore the titer data in the context of specific clades or viral mutations. For example, coloring the H3N2 tree by amino-acid identity at HA site 158 makes it easy to see how some sera have neutralization titers that are markedly affected by mutations at that site (**Supplementary Figure S6**).

## Discussion

We have used a sequencing-based neutralization assay to measure the neutralization of 140 strains covering the current diversity of human influenza A virus by the antibodies in 188 human serum samples taken from individuals of a wide range of ages and geographies. Our results reveal substantial variation in titers both across sera from different individuals and across different viral strains for sera from the same individuals. The heterogeneous antibody landscape captured by our measurements will shape both the distribution of infections and the evolutionary dynamics of seasonal influenza over the coming year, as there is clear evidence that humans are more likely to be infected with strains to which they have low neutralizing antibody titers^3–5,21–23,40^.

Here we have mostly examined the titers after aggregation across sera from many different individuals. Some of the serum-specific patterns in the antibody landscapes are likely shaped by vaccination and infection history^10,12^, and so there is ample room for future analysis of the data in terms of subject age and the available information about recent vaccinations and exposures. Additionally, our study uses sera from individuals of a wide range of ages from several geographic locations (albeit all in the Northern Hermisphere)—but an important area for future work is to determine the role that the antibody immunity of different subsets of the global population play in shaping influenza evolution and transmission^30,40–42^.

The measurements reported here represent the largest-ever single-study dataset of influenza neutralizing antibody titers, with the entire effort being completed in less than six months—with the experimental work performed nearly entirely by a single person. The fact that so much titer data can be generated so rapidly demonstrates how new experimental techniques enable near real-time measurement of the human neutralizing antibody landscape to influenza, potentially enabling the use of such data in immediate public-health applications in addition to retrospective studies.

Here we have made an intentional decision to report the dataset immediately with just simple visualizations, rather than sharing the data only after lengthy post-hoc analyses designed to draw final conclusions. The reason is that these data were generated with the goal of helping inform influenza vaccine-strain selection in September 2025 for the 2026 Southern Hemisphere vaccine; sharing these data immediately facilitates further analysis and interpretation by scientists involved in vaccine-strain selection as well as the larger scientific community. After the completion of this study and posting of the initial version as a *bioRxiv* preprint, these data were used as one of many sources to help inform vaccine strain selection in September 2025^43^. The influenza A components were both updated; the H1N1 component was updated to a D.3.1 strain (Missouri/11/2025 in both egg- and cell-based formulations), while the H3N2 component was updated to a J.2.4 strain (A/Singapore/GP20238/2024 in the egg-based formulation and A/Sydney/1359/2024 in the cell-based formulation)^44^. The H1N1 component strain selected for the vaccine included a 113K mutation that analysis of our large-scale neutralization dataset reported here showed caused measurable antigenic change^43^.

It will only be possible to determine in retrospect the extent to which the data presented here are informative for determining which viral strains or mutations will spread in the human population, or indeed how well the Southern Hemisphere vaccine components will be matched to the circulating variants in 2026. It also remains unclear exactly how the large dataset we report here can best be incorporated into quantitative models^17–20^ of how immune pressure shapes influenza virus evolution, as it is important to remember that antigenic change is only one component of viral fitness. Here we have freely shared the dataset so both ourselves and other scientists can leverage it to advance basic and applied goals related to understanding influenza virus evolution and immunity.

## Acknowledgements

We thank Zoe Munsey and Sophie Rotter-Aboyoun for assistance with collection of the SCH sera. This work was funded in part by the NIH/NIAID under R01AI165281 (to TB and JDB), 75N93021C00015 (to BJC, TB and JDB), F30AI186284 (to CK), R01AI165818 (to SAT and DJS), and 75N93021C00014 (to SAT and DJS). JDB and TB are Investigators of the Howard Hughes Medical Institute. This work was also funded in part by the UK Medical Research Council grant MR/Y004337/1 (to SAT and DJS) and Doctoral Training Grant (SAT). Sera from NIID were provided by Reiko Saito, with support from the Grants-in-Aid for Emerging and Reemerging Infectious Diseases from the Ministry of Health, Labour and Welfare, Japan (grant nos. 24HA2005). This research was also supported by Dolores Covarrubias and the Genomics & Bioinformatics Shared Resource (RRID:SCR_022606) of the Fred Hutch/University of Washington Cancer Consortium (P30 CA015704) and by Fred Hutch Scientific Computing (NIH grants S10-OD-020069 and S10-OD-028685). The EPI-HK study was also funded in part by the Theme-based Research Scheme under project no. T11-712/19-N (to BJC and NHLL) from the Research Grants Council from the University Grants Committee of Hong Kong. We gratefully acknowledge the authors and originating and submitting laboratories of the sequences from the GISAID EpiFlu Database^45^ on which this research is partly based. This manuscript is the result of funding in part by the National Institutes of Health (NIH). It is subject to the NIH Public Access Policy. Through acceptance of this federal funding, NIH has been given a right to make this manuscript publicly available in PubMed Central upon the Official Date of Publication, as defined by NIH.

## Author contributions

Conceptualization: CK, JH, ANL, TB, JDB

Library design: CK, JH, ANL, ST, DJS, JDB

Experiments: CK, ANL

Data analysis: CK, JH, JL, TB, JDB

Collection of sera: IGB, BJC, JAG, ALG, RH, HH, FH, KL, NHLL, NSL, HP, SW

Writing - original draft: CK, JH, TB, JDB

Writing - review and editing: all authors

## Methods

### Data and code availability

See the GitHub repository at https://github.com/jbloomlab/flu-seqneut-2025 for all data and analysis code, including final neutralization titers, barcode counts of each variant in each experiment, plots of all individual neutralization curves, and detailed serum metadata. That GitHub repository includes all details; key summary files are as follows:

- Information on the tested human sera: **Supplementary File 1** and https://github.com/jbloomlab/flu-seqneut-2025/blob/main/results/aggregated_analyses/human_se ra_metadata.csv
- Information on the tested virus strains: **Supplementary File 2** and https://github.com/jbloomlab/flu-seqneut-2025/blob/main/data/viral_libraries/flu-seqneut-2025-ba rcode-to-strain_actual.csv
- All measured neutralization titers after quality-control: **Supplementary 3** and https://github.com/jbloomlab/flu-seqneut-2025/blob/main/results/aggregated_analyses/human_se ra_titers.csv
- Summary statistics on virus-specific titers: https://github.com/jbloomlab/flu-seqneut-2025/blob/main/results/aggregated_analyses/human_se ra_titers_summarized.csv
- Interactive page with links to the all neutralization curves and notebooks showing per-plate and per-serum quality-control: https://jbloomlab.github.io/flu-seqneut-2025/

### Biosafety

All experiments utilized influenza virions with HA ectodomain proteins matching those from recent naturally occurring human seasonal H3N2 or H1N1 influenza strains. Such strains are classified as biosafety-level-2 according to the CDC BMBL handbook (edition 6). The non-HA genes were derived from the lab adapted A/WSN/1933 (H1N1) strain, which is also classified as biosafety-level-2 according to the CDC BMBL handbook. All experimental work involving the viruses or human sera were performed at biosafety-level-2.

### Human sera

Serum samples were sourced from individuals across ages and geographical locations through a combination of residual blood draws from hospitals, epidemiological studies, and vaccination cohorts.

Deidentified pediatric sera were obtained from children who were not immunocompromised and were undergoing routine medical care at Seattle Children’s Hospital (SCH) in April 2025 with approval from the SCH Institutional Review Board with a waiver of consent.

The deidentified remnant sera from University of Washington Medical Center (UWMC) were obtained from individuals testing positive for HBsAb (to control for intact immunity) in March 2025 and was approved by the UW Institutional Review Board with a consent waiver.

The EPI-HK sera were taken from the “<u>E</u>valuating <u>P</u>opulation Immunity in <u>H</u>ong <u>K</u>ong” study, a community-based longitudinal observational cohort study^46^ of approximately 2000 participants of all ages run by the University of Hong Kong since 2020, and for this analysis a subset of 42 cross-sectional sera meant to be representative of this population-based cohort at the end of the local 2024/25 Northern Hemisphere winter influenza season were selected by randomly selecting 3 sera from each 5-year age band between 10-79 years of age collected in March-April 2025. The EPI-HK study protocol was approved by the Institutional Review Board of the University of Hong Kong and written informed consent was obtained from all study participants or their legal guardians.

The sera from NIID were from participants in a vaccine study by the National Institute of Infectious Diseases in Japan; and included 48 sera taken from 48 unique individuals pre-vaccination (October-November 2024) and also 7 sera taken from 7 of these same individuals post-vaccination (November-December 2024) timepoints.

All sera were treated with receptor-destroying enzyme and heat-inactivated prior to use in neutralization assays as described previously^26,31^ in order to eliminate both sialic-acid containing compounds that might non-specifically inhibit viral infection. Briefly, lyophilized receptor-destroying enzyme II (Seikan) was resuspended in 20 mL PBS and vacuum-filtered through a 0.22 uM filter. Then, 25 uL of sera was incubated with 75 uL of receptor-destroying enzyme (constituting a 1:4 dilution) at 37°C for 2.5 hours and then 55°C for 30 minutes. Sera were used immediately or stored at −80°C.

### Design of sequencing-based neutralization assay library

The goal of our library design was to choose HAs from sequenced human strains that were either at high frequency or had mutations that we deemed likely to be of antigenic or evolutionary significance. To select these 2025-circulating strains that we hoped would be representative of future HA diversity, we identified human seasonal H1N1 and H3N2 haplotypes that had been sequenced within a 6-month time period of library design (May 2025), pared down the list of recent haplotypes to only those that had a high local branching index^47^ on Nextstrain^38^ 6-month builds and/or contained mutations at antigenic sites. For H3N2 strains, we so used an analysis approach conceptually similar to that previously described for SARS-CoV-2^48^ to identify HA mutations that have recently independently arisen more than expected from the underlying mutation rate and so are putatively beneficial to the virus, and also selected representative strains containing such mutations. Altogether, this process selected 77 H3N2 strains and 39 H1N1 strains. The code for choosing strains is available at https://github.com/jbloomlab/flu-seqneut-2025/tree/main/non-pipeline_analyses/library_design and the nucleotide and protein sequences for the HA ectodomain for all 2025-circulating strains and recent vaccine strains in the final library are available at https://github.com/jbloomlab/flu-seqneut-2025/tree/main/results/viral_strain_seqs.

In addition, our libraries also included the HAs from the component strains of both H3N2 and H1N1 seasonal vaccine strains. For H3N2, we included cell- and egg-based vaccine strains from the 2014 vaccine to present. For H1N1, we included cell- and egg-passaged vaccine strains from the 2010 vaccine to present. For both H3N2 and H1N1 vaccine strains, some egg-passaged strains had to be dropped as they did not grow to high titers in our system which generates viruses via mammalian cell lines (293T and MDCK-SIAT1-TMPRSS2 cells).

Overall, our final libraries after library generation quality control steps contained 286 barcodes covering 76 recent H3N2 strains, 38 recent H1N1 strains, 19 past H3N2 egg- or cell-produced vaccine strains, and 7 past H1N1 egg- or cell-produced vaccine strains (https://github.com/jbloomlab/flu-seqneut-2025/blob/main/data/viral_libraries/flu-seqneut-2025-barcode-to-strain_actual.csv). Note that the original designed libraries contained slightly more strains and barcodes as a few dropped out during library generation and quality control: the designed libraries included 322 barcodes covering 76 recent H3N2 strains, 39 recent H1N1 strains, 22 past H3N2 egg- or cell-produced vaccine strains, and 9 past H1N1 egg- or cell-produced vaccine strains (https://github.com/jbloomlab/flu-seqneut-2025/blob/main/data/viral_libraries/flu-seqneut-2025-barcode-to-strain_designed.csv).

### Cloning of barcoded HAs

As described previously^26,27,30^, the HA genes used for our libraries consist of the noncoding regions from the lab-adapted A/WSN/1933 (H1N1) HA, the first 19 (for H3 constructs) or 20 (for H1 constructs) amino acids of the N-terminal signal peptide from the A/WSN/1933 HA, the HA ectodomain from each strain of interest, the consensus H3 transmembrane domain (for H3 constructs) or the A/WSN/1933 transmembrane domain (for H1 constructs), the cytoplasmic tail from A/WSN/1933 with synonymous recoding, a double stop codon after the end of the coding sequence, followed by a 16-nucleotide barcode, the Illumina Read 1 priming sequence, and a duplicated packaging signal from A/WSN/1933. This construct enables barcoding of HAs without disrupting viral genome packaging and provides common priming sequences that can be used for barcode amplification.

For cloning the barcoded HAs into influenza reverse genetics plasmids, we used a simplified cloning strategy as described previously^26,27^. Briefly, barcoded constructs encoding the HA ectodomains were ordered from Twist Biosciences with sequence homology at the 5’ end of the coding sequence with the lab-adapted A/WSN/1933 (H1N1) HA signal peptide and sequence homology at the 3’-end (after the barcode) with the Illumina Read 1 sequence. We tagged each HA variant with randomly generated 16-nucleotide barcodes, specifically avoiding barcode sequences used in prior libraries^26,27^ or beginning with GG nucleotides (we found such barcodes can sequence poorly in the sequence context of our libraries). Barcodes fragments were then assembled into plasmid backbones using HiFi Assembly Mastermix (NEB) per manufacturer’s instructions.

The plasmid backbone for both the H1N1 and H3N2 constructs was the derivative of the pHH21 uni-directional reverse genetics plasmid^49^ that we have described previously^26,30^; see https://github.com/dms-vep/flu_h3_hk19_dms/blob/main/library_design/plasmid_maps/2851_pHH_WSNHAflank_GFP_H3-recipient_duppac-stop.gb for a map of this plasmid backbone. This plasmid was digested with enzymes XbaI and BsmBIv2 (NEB) per manufacturer’s instructions and the desired fragment was obtained by gel electrophoresis and purification. Each barcoded construct was then transformed into competent cells in the Bloom lab, and individual colonies were then screened and DNA prepped by Azenta/Genewiz. Exemplar plasmid maps for a H1 and H3 strain are at https://github.com/jbloomlab/flu-seqneut-2025/blob/main/non-pipeline_analyses/library_design/plasmids/example_constructs and the full set of all plasmid maps is at https://github.com/jbloomlab/flu-seqneut-2025/blob/main/non-pipeline_analyses/library_design/plasmids. The barcodes linked to each strain in the final library are at https://github.com/jbloomlab/flu-seqneut-2025/blob/main/data/viral_libraries/flu-seqneut-2025-barcode-to-strain_actual.csv.

### Generation and titering of viruses carrying barcoded HAs

Barcoded viruses expressing the different library HAs were generated using reverse genetics^49,50^. As described previously^26,27^, we generate the two or three barcoded variants for each HA strain in the library by pooling barcoded plasmids encoding that particular HA prior to transfecting cells. However, the barcoded variants for each strain are generated independently. For the reverse genetics, plasmid DNA mixes were made containing 250 ng of a given strain’s HA plasmid pool (containing equal amounts of the two or three of the independently barcoded constructs) with 250 ng of each of a pHW-series bidirectional reverse genetics plasmid^50^ encoding each non-HA segment (PB1, PB2, PA, NA, M, NP, NS) from A/WSN/1933 (H1N1). These DNA mixtures were incubated with 100 uL of DMEM and 3 uL of BioT Transfection Reagent (Bioland Scientific) per manufacturer’s instructions and then transfected onto co-cultures of 5e5 293T cells and 5e4 MDCK-SIAT1-TMPRSS2 cells that had been plated ∼24 hours prior in 6-well dishes in D10 media (DMEM supplemented with 10% heat-inactivated fetal bovine serum, 2 mM L-glutamine, 100 U per mL penicillin and 100 ug per mL streptomycin). At ∼16-20 hours post-transfection, media was removed, cells were washed gently with 2mL of PBS, and then 2 mL of influenza growth media (Opti-MEM supplemented with 0.1% heat-inactivated FBS, 0.3% bovine serum albumin, 100 ug per mL of calcium chloride, 100 U per mL penicillin and 100 ug per mL streptomycin) was added. After an additional ∼45 hour incubation (∼65 hours post transfection), viral supernatants were aliquoted for storage at −80°C and used to set up a single viral passage (intended to reduce carry-over plasmid DNA and increase titers). For these passages, 100 uL of each viral stock was infected onto a single well of a 6-well plate containing 4e5 MDCK-SIAT1-TMPRSS2 cells in 2 mL of influenza growth media for ∼40 hours, as described previously^27^. Supernatants were then cleared of cell debris by centrifugation at 400 g for 5 minutes before being stored at −80°C.

As in prior work^26,27^, to determine the relative transcriptional titer of each of the passaged viruses, we made an equal-volume pool which was serially 2-fold diluted and used to infect MDCK-SIAT1 cells. At 16 hours post infection, cells were lysed and viral barcodes were sequenced as described below. We used these barcode sequencing counts to determine the relative amount of each viral strain (see https://github.com/jbloomlab/flu-seqneut-2025/blob/main/non-pipeline_analyses/library_pooling/notebooks/250716_initial_equal_volume_pool.ipynb), and then repooled so that each strains’ barcodes would be present roughly equally in the pool. This repooled, equally proportioned library stock was then serially 2-fold diluted and used to infect MDCK-SIAT1 cells exactly as described above, and barcode counts were used to affirm roughly equal pooling of strains and to determine the virus dilutions where viral transcription tracked linearly with the amount of virus particles added to cells as described previously^26,27^ (see https://github.com/jbloomlab/flu-seqneut-2025/blob/main/non-pipeline_analyses/library_pooling/notebooks/250723_balanced_repool.ipynb). Based on this analysis, for the experiments described here we chose the 1:16 dilution of the virus library stock as it was in the early part of this linear range where viral transcription is linearly correlated with viral neutralization.

### Sequencing-based neutralization assays

The experimental setup was nearly identical to that outlined in previously^26^ with a few modifications outlined below. A detailed protocol is available on protocols.io at https://dx.doi.org/10.17504/protocols.io.kqdg3xdmpg25/v2. Sera were diluted to 1:20 (accounting for this initial 1:4 dilution from receptor-destroying enzyme treatment) in 50uL of influenza growth media and then serially 2.3-fold diluted down columns of 96-well plate. As determined in virus library titration experiments described above, virus library was then added to all wells at a 1:16 dilution in 50 uL, resulting in a range of serum dilutions from 1:40 to 1:13,619 across each 8-well column of a 96-well plate. These virus serum-mixtures were then incubated at 37°C with 5% CO2 for 1 hour before 1.5e5 MDCK-SIAT1 cells were added per well in a total of 50 uL of influenza growth media. After a 16 hour incubation, cells were lysed and barcodes were sequenced as described previously. Briefly, barcoded RNA spike-in (prepared as described previously^27^) was diluted to 2 pM in iScript Sample Preparation Reagent (BioRad) and incubated on cells for 5 minutes. Lysate was then transferred to new 96-well plates for storage, and 1 uL of lysate was used in 10uL cDNA synthesis reactions using the iScript cDNA Synthesis Kit (BioRad) per manufacturer’s instructions. As described previously^26^, we then amplified cDNA in two rounds of PCR. The first round of PCR added a 6-bp index, allowing us to multiplex different plates using the same dual indices (added in the second round of PCR), which helped decrease sequencing costs. For this first round PCR, forward primers were either one the same four forward primers described previously^26^ or a similar new forward primer (listed below). In all cases, this forward primer was paired with the same reverse primer described previously^27^. For this first round of PCR, we used 2 uL of cDNA template in 25 uL PCR reactions using KOD Polymerase Hot Start 2x Mastermix (Sigma) per manufacturer’s instructions. The four new forward primers were:

- *5’-GTGACTGGAGTTCAGACGTGTGCTCTTCCGATCTgtctaaCCTACAATGTCGGATTTGTATTTAA TAG-3’*
- *5’-GTGACTGGAGTTCAGACGTGTGCTCTTCCGATCTacgctgCCTACAATGTCGGATTTGTATTTAA TAG-3’*
- *5’-GTGACTGGAGTTCAGACGTGTGCTCTTCCGATCTtatagcCCTACAATGTCGGATTTGTATTTAAT AG-3’*
- *5’-GTGACTGGAGTTCAGACGTGTGCTCTTCCGATCTcgagctCCTACAATGTCGGATTTGTATTTAA TAG-3’*)

In the second round of PCR, unique dual indexing primers (described previously^27^) were added in 25 uL reactions, again using KOD Polymerase Hot Start 2x Mastermix (Sigma) per manufacturer’s instructions. These second round PCR products were then pooled at equal volume, gel extracted, purified, quantified and sequenced exactly as described previously^26^.

### Analysis of sequencing data to determine neutralization titers

The sequencing data were analyzed as described previously^26,27^ using the *seqneut-pipeline* (https://github.com/jbloomlab/seqneut-pipeline), version 4.0.1. Briefly, in this pipeline, the Illumina sequencing data are parsed to count each barcoded variant in each well of each plate (see https://github.com/jbloomlab/flu-seqneut-2025/tree/main/results/barcode_counts for these counts). The barcode counts are then normalized to fractional infectivity of each variant at each serum concentration using the counts of the RNA spike-in standard (see the “frac_infectivity.csv” files for each plate at https://github.com/jbloomlab/flu-seqneut-2025/tree/main/results/plates). The *neutcurve* package (https://github.com/jbloomlab/neutcurve) package is used to fit Hill curves for each variant and serum, and the titer is quantified as the midpoint of the curves. See the analysis configuration file (https://github.com/jbloomlab/flu-seqneut-2025/blob/main/config.yml) for details about the parameters used for the curve fitting and subsequent quality-control to remove low-quality curves. For HAs with multiple barcodes, we report the median titer across barcodes.

A fully reproducible *Snakemake*^51^ pipeline that performs the above analysis is available on GitHub at https://github.com/jbloomlab/flu-seqneut-2025. See https://jbloomlab.github.io/flu-seqneut-2025 for HTML rendering of all neutralization curves, quality control notebooks, and interactive plots summarizing the data.

### Phylogenetic analyses with Nextstrain

For each subtype, we created a phylogenetic tree for the HA nucleotide sequences used in the neutralization assays along with approximately 500 additional HA sequences from the Global Initiative on Sharing All Influenza Data (GISAID) EpiFlu database^45^ collected between August 1, 2024 and August 22, 2025 to show the genetic context of library sequences and the genetic diversity that has emerged globally since we finalized the library design. We required contextual sequences to have complete collection dates and we excluded known outliers that we previously identified through weekly genomic surveillance analyses with Nextstrain (H3N2 outliers at https://github.com/nextstrain/seasonal-flu/blob/825314f/config/h3n2/outliers.txt and H1N1pdm outliers at https://github.com/nextstrain/seasonal-flu/blob/825314f/config/h1n1pdm/outliers.txt). We randomly sampled the contextual sequences, evenly sampling from each major global region and month. We aligned sequences to a reference virus sequence (A/Wisconsin/67/2005 for H3N2 and A/Wisconsin/588/2019 and H1N1pdm) with Nextclade version 3.16.0^52^ and inferred a divergence tree with IQ-TREE version 3.0.1^53^ using augur tree^54^. We rooted the tree with the reference virus, pruned the reference virus from the tree, and inferred a time tree with TreeTime version 0.11.4^55^ using a fixed clock rate (0.00382 for H3N2 and 0.00329 for H1N1pdm), a clock standard deviation of 1/5th the clock rate, a constant coalescent, marginal date inference, stochastic polytomy resolution, and FFT inference. We inferred ancestral nucleotide and amino acid sequences for internal nodes with TreeTime through the augur ancestral command and used the mutations associated with these inferred sequences to annotate clades with augur clades. For each tree, we created a corresponding measurements panel^39^ containing the log_2_ neutralization titers per serum id. These analyses are available through the reproducible *Snakemake* pipeline at https://github.com/blab/kikawa-seqneut-2025-VCM/. A complete list of GISAID accessions and authors is available at https://github.com/blab/kikawa-seqneut-2025-VCM/blob/79668f4/gisaid_accessions.tsv.

To infer phylogenetic trees from HA protein sequences such that branch length reflects the number of amino acid mutations separating different viral strain HA sequences, we developed a separate *Snakemake* pipeline and integrated our titer data with this phylogenetic analysis as well. The pipeline for building the trees is described at https://github.com/jbloomlab/nextstrain-prot-titers-tree, with the configuration for our analysis placed at https://github.com/jbloomlab/flu-seqneut-2025/blob/main/config.yml. These protein-based trees can be explored at https://nextstrain.org/community/jbloomlab/flu-seqneut-2025@main/H3N2 and https://nextstrain.org/community/jbloomlab/flu-seqneut-2025@main/H1N1.

## Supplementary Material

**Supplementary Figure S1.**
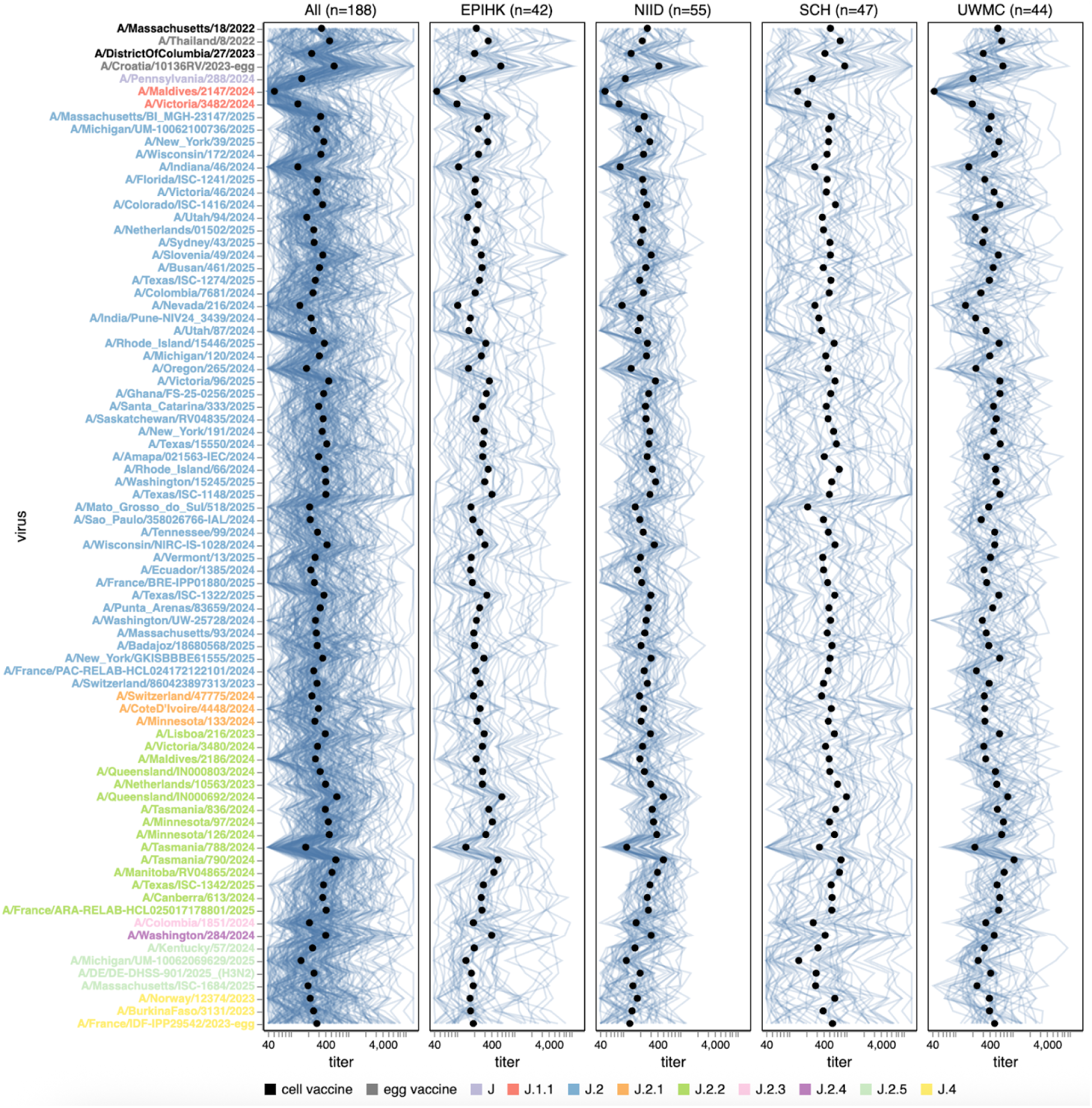
Titers of individual human sera to recent H3N2 strains. Each blue line shows the titers of a serum against the 76 H3N2 strains that capture current circulating diversity, as well as the last two sets of egg- and cell-produced vaccine strains. Black points show the median titer across all sera. Strain labels are colored by whether a strain is a vaccine strain or its subclade per the legend at bottom. The lower limit of detection for titers in our neutralization assays was 40. See https://jbloomlab.github.io/flu-seqneut-2025/human_sera_titers_H3N2_recent_individual_sera.html for an interactive version of this plot that allows mousing over points and lines for details about individual sera or viruses, and subsetting on sera from specific age ranges.

**Supplementary Figure S2.**
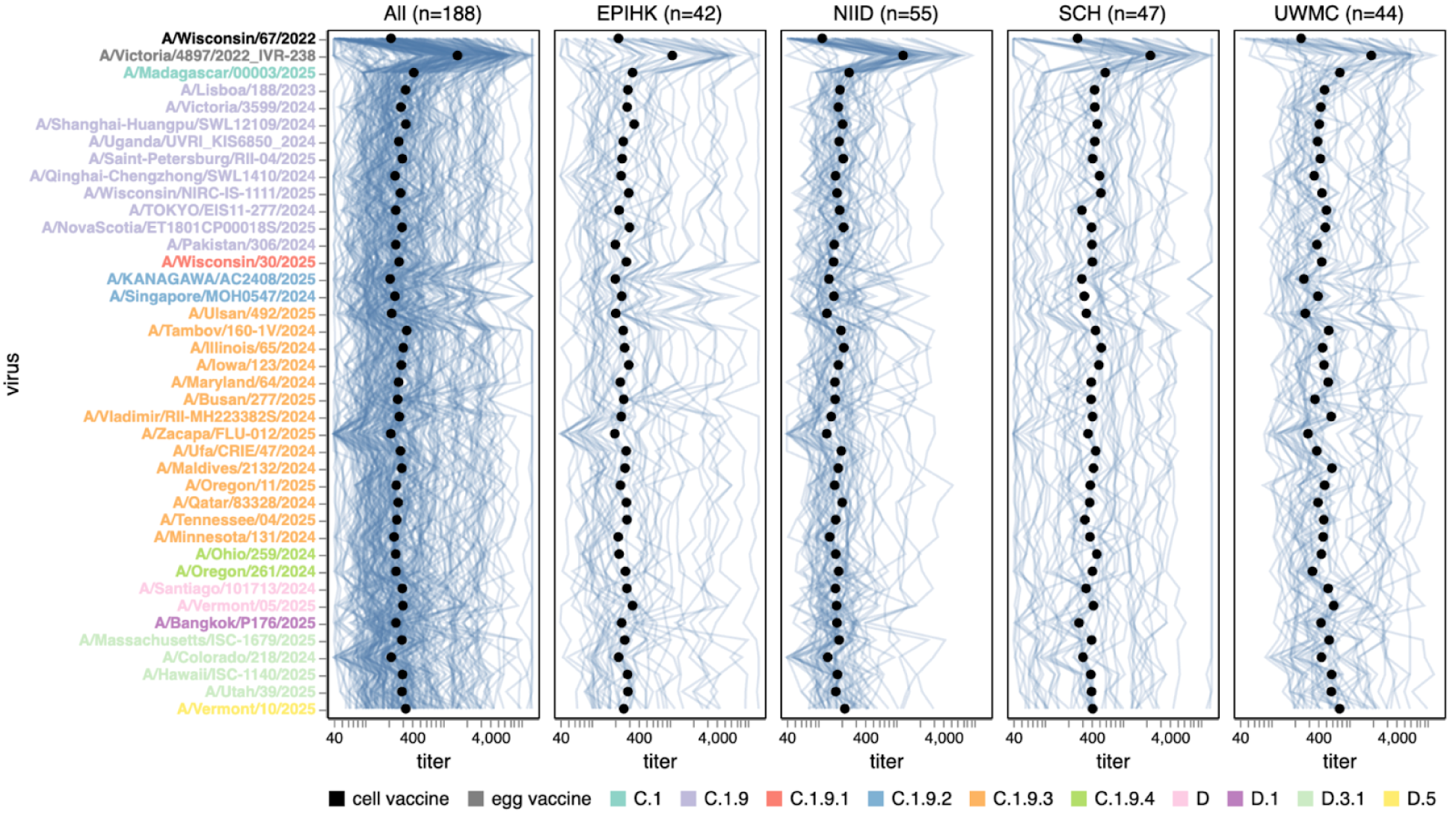
Titers of individual human sera to recent H1N1 strains. Each blue line shows the titers of a different serum against the 38 H1N1 strains that capture current circulating diversity, as well as the last set of cell- and egg-produced vaccine strains. The black points show the median titer across all sera. The leftmost plot shows the titers against all sera, and the remaining plots show titers for sera from each different cohort. Strain labels are colored by whether a strain is a cell- or egg-produced vaccine strain or its subclade per the legend at bottom. The lower limit of detection for titers in our neutralization assays was 40. See https://jbloomlab.github.io/flu-seqneut-2025/human_sera_titers_H1N1_recent_individual_sera.html for an interactive version of this plot that allows mousing over points and lines for details about individual sera or viruses, and subsetting on sera from specific age ranges.

**Supplementary Figure S3.**
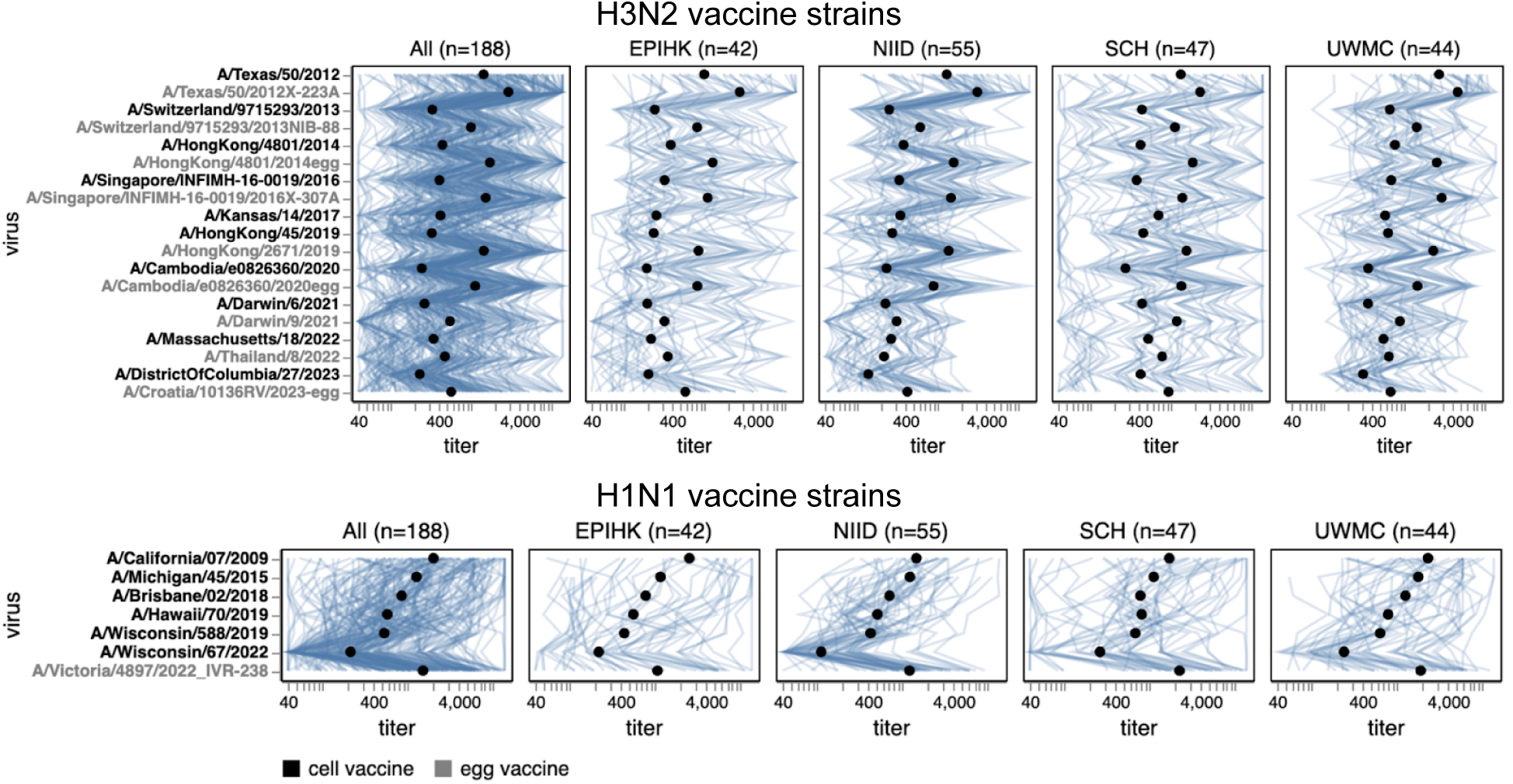
Titers of individual human sera to past vaccine strains. Each blue line shows the titers of a different serum against the viral strains, and the black points show the median titer across all sera. The top panel shows past H3N2 vaccine strains, and the bottom panel shows past H1N1 vaccine strains. The leftmost plot shows the titers against all sera, and the remaining plots show titers for sera from each different cohort. Strain labels are colored by whether a strain is a cell- or egg-produced vaccine strain per the legend at bottom. The lower limit of detection for titers in our neutralization assays was 40. See https://jbloomlab.github.io/flu-seqneut-2025/human_sera_titers_H3N2_vaccine_individual_sera.html and https://jbloomlab.github.io/flu-seqneut-2025/human_sera_titers_H1N1_vaccine_individual_sera.html for an interactive version of this plot that allows mousing over points and lines for details about individual sera or viruses, and subsetting on sera from specific age ranges.

**Supplementary Figure S4.**
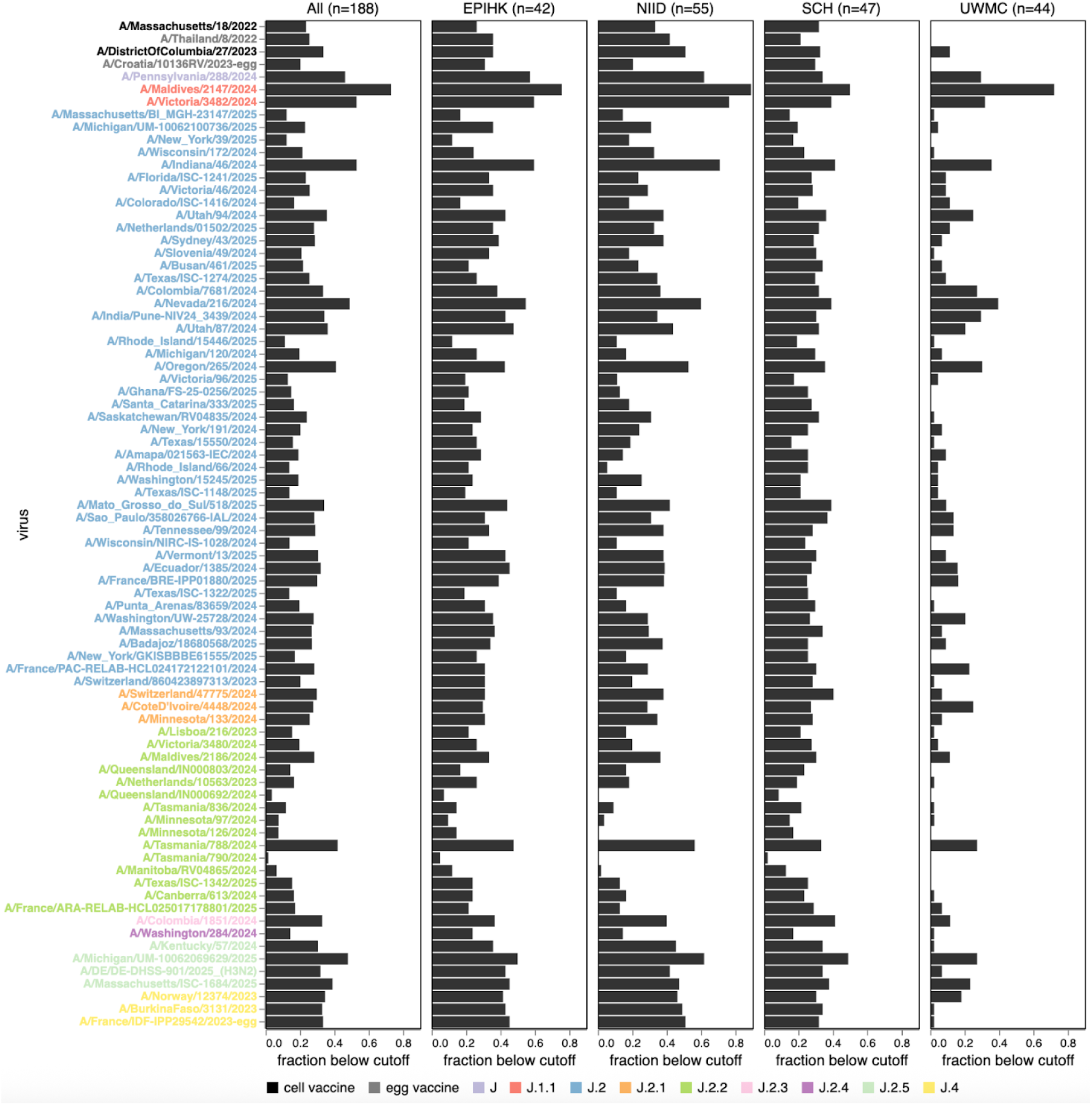
Fraction of sera with low titers to each recent H3N2 strain. Each bar indicates the fraction of sera with a neutralization titer below 140 for each of the 76 H3N2 strains that capture current circulating diversity, as well as the last two sets of egg- and cell-produced vaccine strains. The leftmost panel shows the fractions for all sera, and the other panels show fractions for each subset of sera. Strain labels are colored by whether a strain is a vaccine strain or its subclade per the legend at bottom. See https://jbloomlab.github.io/flu-seqneut-2025/human_sera_titers_H3N2_recent_frac_below_cutoff.html for an interactive version of this figure that enables adjustment of the titer cutoff used to compute the fraction of sera below the cutoff, mousing over points for details, and subsetting on sera from specific age ranges. These values are available as a flat file at https://github.com/jbloomlab/flu-seqneut-2025/blob/main/results/aggregated_analyses/human_sera_titers_summarized.csv for downstream analysis.

**Supplementary Figure S5.**
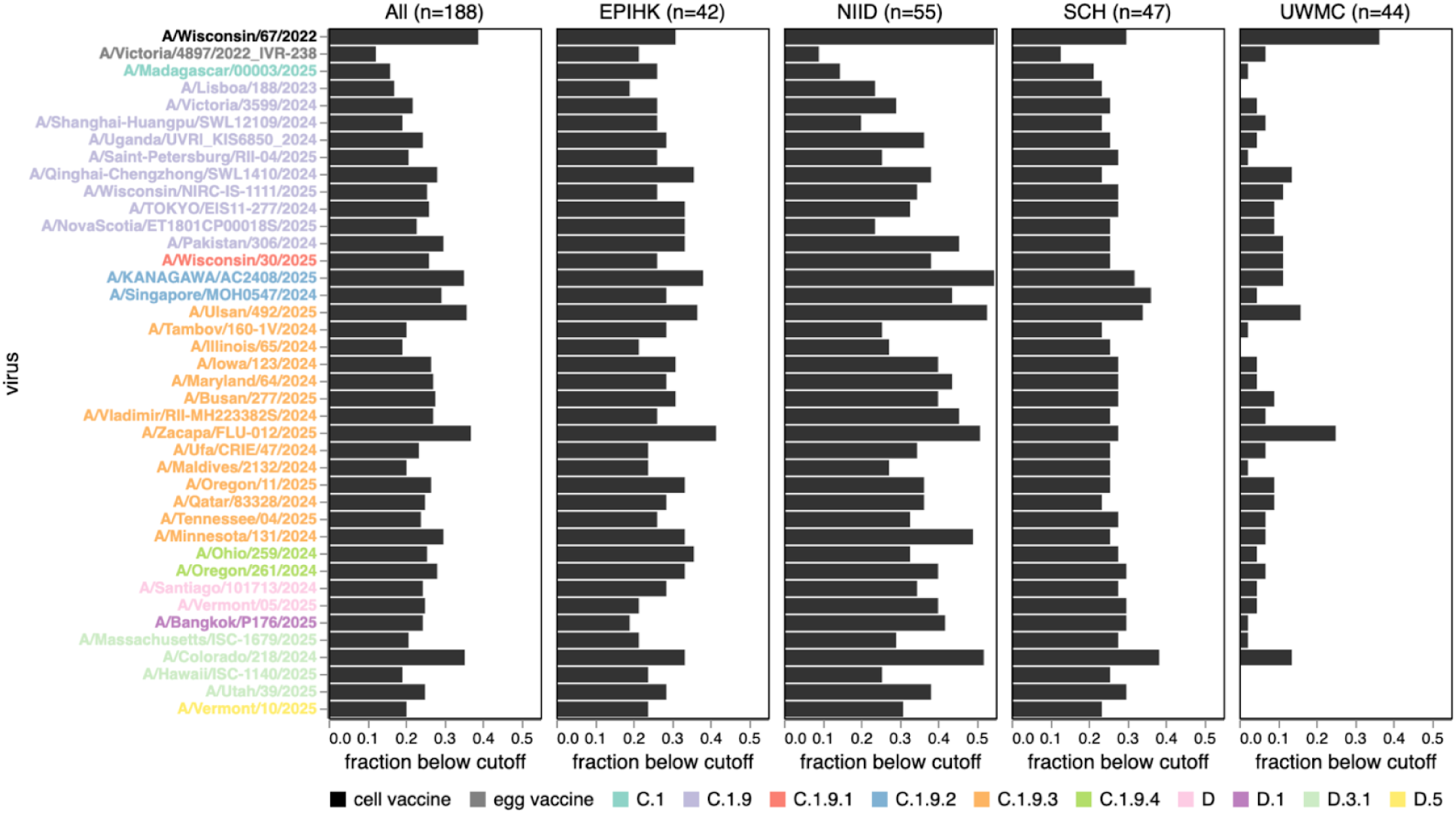
Fraction of sera with low titers to each recent H1N1 strain. Each bar indicates the fraction of sera with a neutralization titer below 140 for each of the 38 H1N1 strains that capture current circulating diversity, as well as the last set of egg- and cell-produced vaccine strains. The leftmost panel shows the fractions for all sera, and the other panels show fractions for each subset of sera. Strain labels are colored by whether a strain is a vaccine strain or its subclade per the legend at bottom. See https://jbloomlab.github.io/flu-seqneut-2025/human_sera_titers_H1N1_recent_frac_below_cutoff.html for an interactive version of this figure that enables adjustment of the titer cutoff used to compute the fraction of sera below the cutoff, mousing over points for details, and subsetting on sera from specific age ranges. These values are available as a flatfile at https://github.com/jbloomlab/flu-seqneut-2025/blob/main/results/aggregated_analyses/human_sera_titers_summarized.csv for downstream analysis.

**Supplementary Figure S6.**
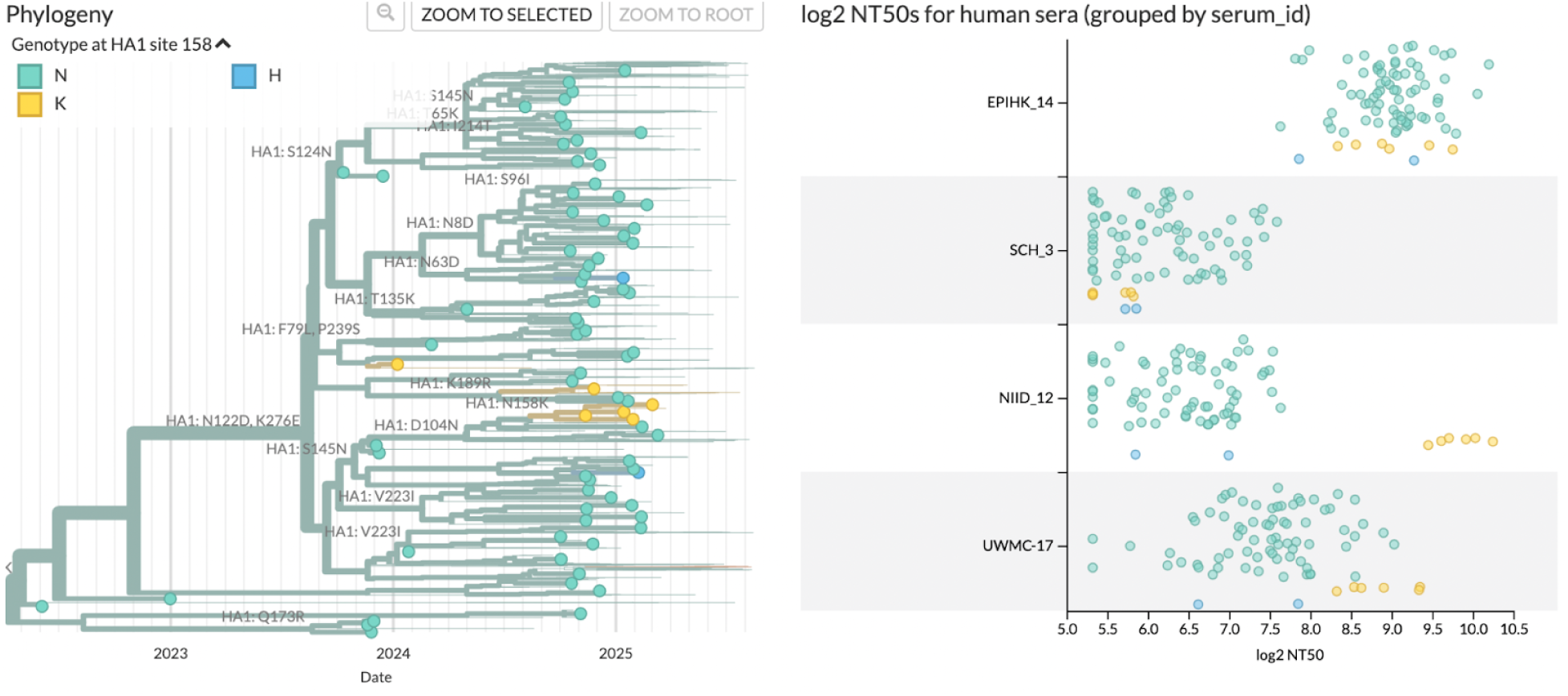
Screenshot of Nextstrain interactive visualization showing the H3N2 HA phylogenetic tree (left) and log_2_ titer measurements for four serum samples (right) colored by amino-acid identity at HA1 position 158. This figure demonstrates how the interactive trees can be used to examine how specific mutations affect titers. The phylogeny (identical to that shown in Figure 2 and Figure 8) is filtered to show circles at tip labels only viruses with neutralization titers. The measurements panel shows the variability of titers between and within serum samples with respect to the amino-acid identity at site 158. For example, samples NIID_12 and UWMC-17 have high titers against all 158K viruses, while sample SCH_3 has low titers against 158K and 158H viruses. See https://nextstrain.org/groups/blab/kikawa-seqneut-2025-VCM/h3n2?branchLabel=aa&c=gt-HA1_158&d=tree,measurements&f_kikawa=present_1&focus=selected&gmax=1052&gmin=72&mf_serum_id=EPIHK_14&mf_serum_id=SCH_3&mf_serum_id=NIID_12&mf_serum_id=UWMC-17&p=grid&tl=none for an interactive view of the tree shown here; note that this interactive tree allows similar visualizations for arbitrary mutations or sera.

**Supplementary File 1. CSV with details about all sera assayed in the experiments.**

This CSV gives details about all sera assayed in the experiments reported here. See https://github.com/jbloomlab/flu-seqneut-2025/blob/main/results/aggregated_analyses/human_sera_metadata.csv for a copy of this CSV.

**Supplementary File 2. CSV with details of all strains and barcodes in the library.**

This CSV gives the barcode, strain name, nucleotide and protein HA ectodomain sequence, and other details about all viral strains in the library. See https://github.com/jbloomlab/flu-seqneut-2025/blob/main/data/viral_libraries/flu-seqneut-2025-barcode-to-strain_actual.csv for a copy of this CSV.

**Supplementary File 3. CSV with all titers measured in the experiments reported here.**

This CSV gives the neutralization titers measured in the experiments reported here, averaged across barcodes and replicates for each strain. See https://github.com/jbloomlab/flu-seqneut-2025/blob/main/results/aggregated_analyses/human_sera_titers.csv for a copy of this CSV.

